# Nanoparticle Accumulation and Penetration in 3D Tumor Models: the Effect of Size, Shape, and Surface Charge

**DOI:** 10.1101/2024.12.06.626955

**Authors:** Pierre Cybulski, Maria Bravo, Jim Jui-Kai Chen, Indra Van Zundert, Sandra Krzyzowska, Farsai Taemaitree, Hiroshi Uji-i, Johan Hofkens, Susana Rocha, Beatrice Fortuni

## Abstract

Preclinical studies have demonstrated that nanoparticles (NPs) hold significant potential for advancing cancer therapy by enhancing therapeutic efficacy while reducing side effects. Their effectiveness in solid tumors is, however, often constrained by insufficient accumulation and penetration. Understanding how the physicochemical properties of NPs – such as size, shape, and surface charge – influence their interaction with cells within the tumor is critical for optimizing NP design. In this study, we addressed the challenge of inconsistent NP behavior by systematically evaluating NP uptake in both 2D and 3D tumor models, and NP penetration in spheroids. Our results showed that larger NPs exhibited higher internalization rates in 2D models but limited penetration in 3D spheroids. Furthermore, negatively charged NPs consistently achieved superior accumulation and deeper penetration than neutral and positively charged NPs. Spherical NPs outperformed rod-shaped NPs in tumor accumulation and penetration. These findings underscore the importance of carefully tailoring NP properties to the complex tumor microenvironment for improved therapeutic outcomes in real tumors.

## Introduction

In recent decades, cancer research has increasingly prioritized developing more efficient and less invasive treatments, as cancer remains a leading cause of death worldwide.^1^ Nanoparticle (NP)-based therapies have emerged as a promising approach, offering improved tumor targeting and accumulation while reducing side effects.^2^ However, despite extensive research and numerous studies demonstrating their efficacy in cancer cells and animal models, clinical adoption remains limited. By 2023, only 14 cancer nanomedicines have been approved by the Food and Drug Administration (FDA), despite more than 25,000 publications on the topic.^3,4^ This disparity underscores the challenges in translating NP-based therapies from the lab to the clinical practice.

Preclinical testing is predominantly based on 2D cell models, which often fail to predict *in vivo* outcomes.^5^ Results from 2D studies are generally not reproducible *in vivo* due to significant biological differences between cell monolayers and animal models.^6^ One key difference is the absence of the extracellular matrix (ECM) in 2D cultures – a critical component of the tumor microenvironment. The dense and complex structure of ECM, with high fluid pressure and low pH, can hinder NP accumulation and penetration, reducing therapeutic efficacy.^7,8^ To address these issues, more biologically relevant 3D cell models, such as multicellular tumor spheroids, have been introduced for testing nanomedicines.^6^

A key challenge in improving NP-based therapies efficacy in solid tumors is understanding how physicochemical properties, such as size, shape, and surface charge, influence their cellular uptake and tumor penetration.^9^ Although these properties have been extensively studied in 2D cell monolayers,^10,11^ research using 3D models like tumor spheroids remains limited and often inconsistent. Identifying the optimal NP properties for better penetration and accumulation in 3D models is crucial to boost clinical translatability of NP-based therapies.

A few studies have explored the optimal NP size for deeper spheroid penetration across a range of sizes. Huang’s work revealed that 2 nm NPs accumulated and penetrated more effectively than 6 nm and 15 nm particles.^12^ Pratiwi and Tchoryk reported that NPs in the 25-50 nm range exhibited similar accumulation and penetration, outperforming larger 100-200 nm NPs.^13,14^ On the other hand, Agarwal observed no significant difference in accumulation or penetration between 100, 200, and 500 nm spheres.^15^ Collectively, these reports suggest smaller NPs tend to achieve deeper penetration in 3D spheroids, though the ideal size remains unclear. The effect of surface charge adds further complexity, with studies often presenting contradictory results. For instance, Wang et al. reported enhanced penetration for positively charged NPs in both 3D spheroids and *in vivo* tumors,^16^ while Sujai et al., in a later study, found that negative gold NPs (AuNPs) accumulated more and penetrated deeper in similar models.^17^ Expanding these challenges, the few studies on NP shape report conflicting results. Zhang et al. showed that elongated NPs, such as nanorods, penetrated spheroids more effectively than spherical ones^18^, while Zhao et al. observed better penetration depth with spherical NPs over rods.^19^ The inconsistencies underscore the challenges in defining optimal NP properties, highlighting the need for a systematic approach to understand how size, charge, and shape influence the behavior of NPs both in 2D and in 3D environments.

This study aims to shed light on the impact of size, surface charge, and shape on NP behavior, in 2D cell monolayers and 3D spheroids. A major challenge in such comparative studies is to isolate the impact of each parameter while keeping others constant. For this purpose, AuNPs of different sizes, shapes, and surface charges were incubated with A549 lung carcinoma cells, chosen for their ability to form compact, tumor-like spheroids, closely mimicking solid tumors. Confocal fluorescence microscopy was used to monitor NP internalization in 2D models, and NP accumulation and penetration in 3D models. AuNPs were chosen as a model due to their highly tunable size, shape and surface chemistry.^20^ Moreover, their intrinsic photoluminescence (PL) allows for precise and fluorophore-free imaging under two-photon excitation.^21^ To accurately assess the penetration of each NP type in 3D models, we used an in-house developed algorithm for analysis of the fluorescence images acquired.

These findings will be valuable for optimizing NP designs that better address the complexities of tumor microenvironments, potentially enhancing the efficacy of NP-based therapies. By understanding how specific NP properties impact accumulation and penetration, this study lays the groundwork for tailoring NP features to achieve improved therapeutic outcomes.

## Materials and methods

### Materials

AuNPs were acquired from Nanopartz Inc, with part numbers A11-15-CIT-DIH-1-25 (AuNS10), A11-35-CIT-DIH-1-25 (AuNS30), A11-55-CIT-DIH-1-25 (AuNS50), A11-75-CIT-DIH-1-25 (AuNS65), CC12-10-750-NEG-DIH-50-1 [AuNR(-)], 10-750-ZERO-DIH-50-1 [(AuNRØ)], and 10- 750-POS-DIH-50-1 [AuNR(+)]. Dulbecco’s Modified Eagle Medium (DMEM), gentamicin, phosphate buffered saline (PBS), formaldehyde, trypsin-EDTA (0.5%), Hank’s balanced salt solution (HBSS), Triton X-100 (0.1%), CellMask™ Green and Deep Red were obtained from ThermoFisher Scientific. Fetal bovine serum and Glutamax were purchased from Sigma Aldrich. Phalloidin CruzFluor™ 488 and 647 were obtained from Santa Cruz Biotechnology.

### Cell Culture

A549 cells were maintained in T25 culture flasks at 37°C in a 5% CO□ atmosphere. The cells were grown in cell culture medium (DMEM supplemented with 10% FBS, 1% Glutamax, and 0.1% gentamicin), and were passaged using trypsin-EDTA when a confluency of 80-90% was reached.

### Cell monolayer assays (2D)

A549 cells were seeded in 29 mm glass-bottom dishes (D29-14-1.5-N, CellVis) and grown to 60% confluency. NPs were added at ∼8×10^1^□ NPs/mL for size and shape studies, and ∼4×10^11^ NPs/mL for charge studies. After 3 h of incubation, cells were washed three times with PBS, fresh culture medium was added and the samples were incubated for additional 24 h at 37°C. Before imaging, the samples were stained with CellMask™ Deep Red (AuNSs) or CellMask™ Green (AuNRs – 1 μM in HBSS, 5 min), washed three times with PBS, and imaged in HBSS.

### Spheroid assays (3D)

At 90% confluency, A549 cells were harvested for spheroid formation. Spheroids were formed in agarose microtissues with 256 wells (16×16 array), cast from ultra-pure agarose (20 mg/mL in Milli-Q water with 9 mg/mL NaCl) using a 3D micro-mold (3D Petri Dish^®^, Sigma Aldrich). Once solidified, the microtissue was placed in a culture plate well and equilibrated with culture medium for 15 min. A 190 μL of a cell suspension (∼2.5×10^5^ cells) was added, allowing the cells to settle for 10 min before adding additional culture medium around the microtissue.

After 4 days of incubation, spheroids with diameters of 250 to 350 μm were formed and the medium surrounding the microtissues was carefully removed. 190 μL of NP-containing medium was added to the wells (∼8×10^9^ NPs/mL for size; ∼4×10^11^ NPs/mL for charge; ∼5×10^1^□ NPs/mL for shape). After 10 min, more medium was added around the microtissues to avoid diluting the NP-containing medium inside. Spheroids were incubated with NPs for 24 h, then the medium was removed, and 1 mL of warm PBS was pipetted over the microtissues to dislodge spheroids, which were then collected and transferred to 1.5 mL microcentrifuge tubes.

Harvested spheroids were stained as previously reported.^22^ Briefly, they were washed three times with PBS, centrifuged at 2000 g for 10 s between washes, then fixed in 500 uL of 4% paraformaldehyde for 20 min, followed by another three PBS washes. Permeabilization was done using 1 mL of 0.1% Triton X-100 in PBS for 20 min. Spheroids were then incubated with 200 μL of Phalloidin CruzFluor™ 488 (AuNR) or Phalloidin CruzFluor™ 647 (AuNS) in 3% BSA at a 1:1000 dilution overnight at room temperature. The next day, spheroids were washed three times with PBS and prepared for imaging.

### Fluorescence microscopy

Live-cell imaging was performed in HBSS at 37°C and 5% CO□, using a Leica TCS SP8 DIVE inverted microscope equipped with a 63× oil immersion objective (NA 1.4) for 2D models and a 25× water immersion objective (NA 0.95, FLUOTAR VISIR) for 3D models. A multiphoton InSight X3 laser (Spectra-Physics®) was used for AuNPs imaging (AuNS: 800 nm, 3.8 mW at the objective, 525–575 nm detection; AuNRs: 1300 nm, 3.8 mW, 655-755 nm detection. Diode lasers were used for the fluorescence probes used: 488 nm diode laser for CellMask™ Green and Phalloidin CruzFluor™ 488 (detection: 491-541 nm); 638 nm diode laser for CellMask™ Deep Red and Phalloidin CruzFluor™ 647 (detection: 650–700 nm). Sequential scanning was used to avoid crosstalk between the channels. Image stacks of 1024×1024 pixels were acquired at a scanning speed of 400 Hz, with three-line averaging and a 0.3 or 1 µm z-step, for 2D or 3D assays, respectively.

### Spheroid penetration analysis

At least 15 spheroids per condition were analyzed using an in-house algorithm to calculate the mean NP photoluminescence (PL) intensity across penetration depth. The algorithm first plots the spheroid diameter along the z-axis, allowing the user to select the z range with the largest diameter for analysis. Each xy image in the chosen z range is segmented into 5 µm rings and the mean NP PL is calculated for each ring (snapshots of the software in Figure S13). The values from each stack were normalized (min-max) to generate a penetration curve of each spheroid and the mean penetration was calculated by averaging the normalized curves across all spheroids in each condition.

## Results

### Nanoparticle characterization

Before testing the different AuNPs in cell models, their physicochemical properties were characterized (Figure 1). Electron microscopy (EM) images showed that gold nanospheres (AuNSs) had a uniform spherical shape, with mean diameters of 12, 31, 49, and 64 nm, corresponding to AuNS10, AuNS30, AuNS50, and AuNS65, respectively (Figure 1A-D, F). For the charge effect investigation, gold nanorods (AuNRs) with negative, neutral, and positive surface modifications were selected and named AuNR(-), AuNRØ, and AuNR(+). Transmission EM confirmed their rod-shaped structure, with dimensions of ∼50 nm by ∼15 nm (Figure 1E, size distributions in Figure S1C).

**Figure 1.**
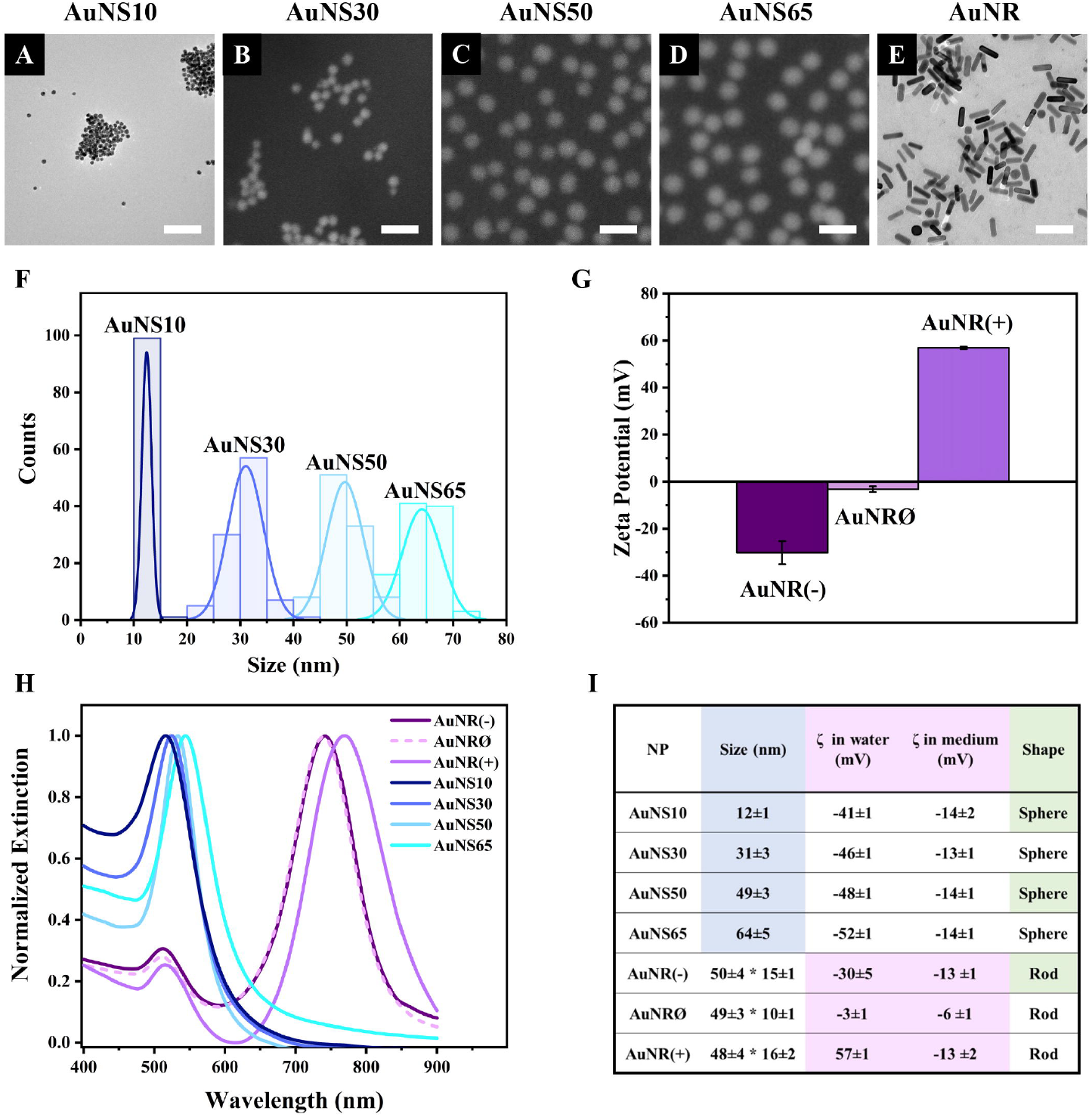
Nanoparticle characterization. Electron microscopy images of **(A)** AuNS10, **(B)** AuNS30, **(C)** AuNS50, **(D)** AuNS65, **(E)** AuNR(-); since no differences were observed between the AuNR samples in EM images, AuNR(-) is shown as a representative, while AuNRØ and AuNR(+) are displayed in Figure S1A-B. Scale bars: 100 nm. **(F)** AuNSs size distribution based on electron microscopy images. **(G)** Zeta potential of AuNR(-), AuNRØ, and AuNR(+). **(H)** Extinction spectra in water. **(I)** Summary table with values shown as mean ± standard deviation.

For all the experiments here reported, NPs were stored as water-based colloidal solutions and small volumes of these solutions were added to the cell culture medium for NP incubation with cells. The surface charges of the different NPs were therefore estimated both in water and in cell culture medium using Zeta potential measurements. The results are summarized in Figure 1I. In water, the Zeta potentials of AuNR(-), AuNRØ, and AuNR(+) were -30 mV, -3 mV, and 57 mV, respectively (Figure 1G). Zeta potentials of the different spherical NPs were consistent across the different sizes: - 41 mV for AuNS10; -46 mV for AuNS30; -48 mV for AuNS50; and -52 mV for AuNS65 (Figure S1D). While suspension in cell medium did not significantly alter the surface charge of the AuNSs, AuNR(-) and AuNRØ, it caused a major change in AuNR(+), reducing the Zeta potential from 57 mV in water to -13 mV (Figure 1I).

The optical properties of each AuNP were characterized by measuring the localized surface plasmon resonance (LSPR) bands using UV-VIS spectroscopy (Figure 1H). The extinction spectra of AuNS10, AuNS30, AuNS50, and AuNS65 exhibited a single plasmonic band at 516 nm, 524 nm, 533 nm, and 544 nm, respectively. A red shift of the plasmonic peak normally occurs with the increase in NP size.^23^ In contrast, AuNRs showed two LSPR peaks – transverse and longitudinal – typical of rod-shaped AuNPs.^24^ These peaks were at 512 nm and 741 nm for AuNR(-), 511 nm and 739 nm for AuNRØ, and 515 nm and 769 nm for AuNR(+). The slight shifts in peak positions among the three AuNR samples are attributed to their different surface modifications,^25^ which account for the different surface charges.

### Influence of NP size

To study the effect of NP size in 2D cell models, A549 human lung adenocarcinoma cells were cultured in 2D monolayers and incubated for 3 h with AuNS10, AuNS30, AuNS50, and AuNS65. Figures 2A-D show representative xy planes from z-stacks, alongside xz and yz orthogonal views, for each NP size. AuNS10 exhibited low and heterogeneous internalization (Figure 2A and S2), while a significant increase in uptake was observed for AuNS30, AuNS50, and AuNS65 (Figures 2B-D and S2). Among these, AuNS65 had the highest uptake, followed by AuNS30 and AuNS50.

**Figure 2.**
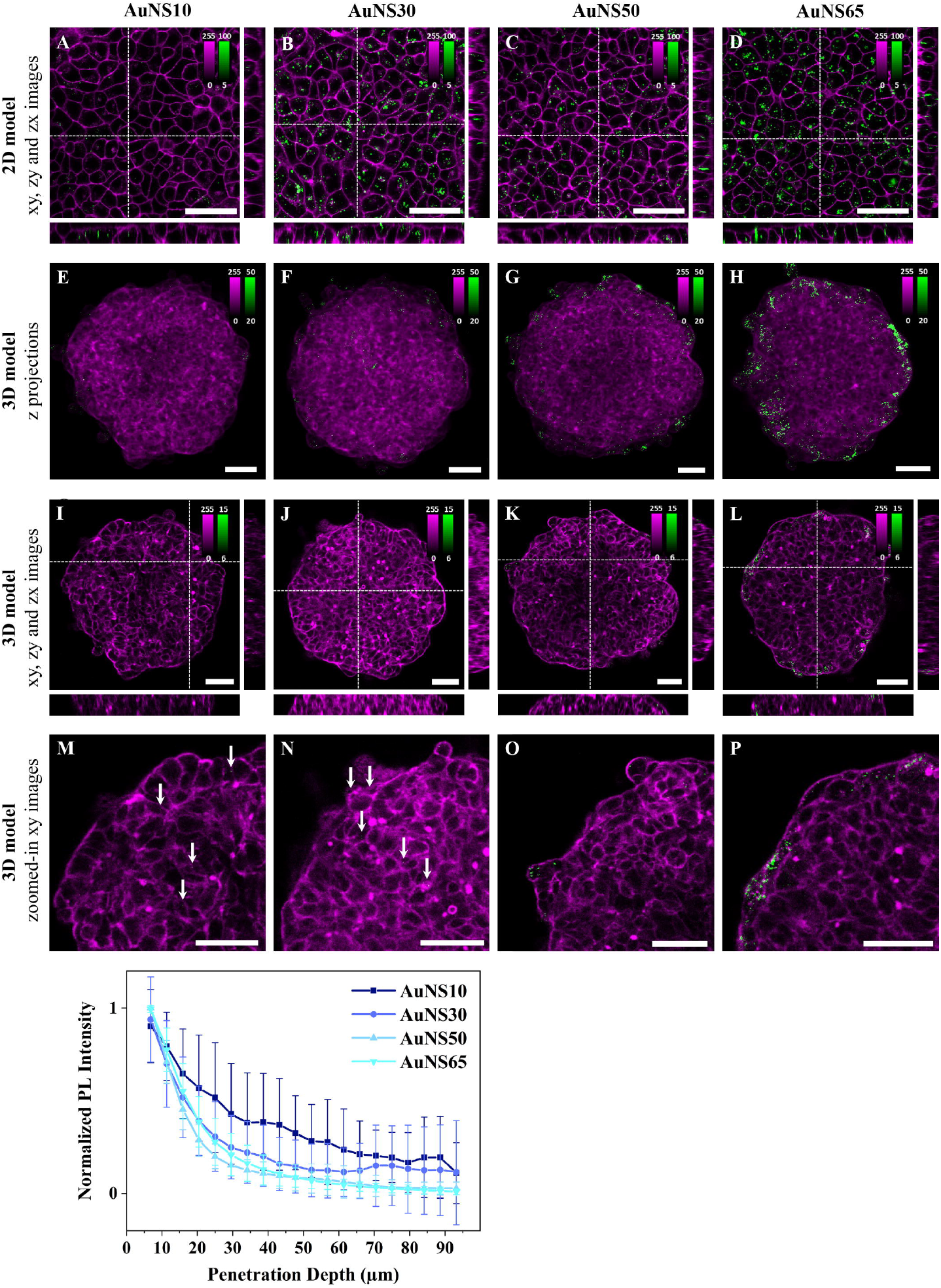
The effect of NP size on cellular uptake and spheroid penetration. Representative confocal fluorescence images of AuNS10, AuNS30, AuNS50, and AuNS65 in **(A-D)** 2D monolayers and **(E-P)** 3D spheroids of A549 cells. Cells were incubated with ∼8×10^1^□ NPs/mL for 3 h or ∼8×10^9^ NPs/mL for 24 h, for 2D or 3D models, respectively. NP PL is shown in green while cellular staining appears in magenta (CellMask^TM^ Deep Red plasma membrane staining for 2D models and Phalloidin CruzFluor^TM^ 647 for cytoskeleton staining in 3D models). Scale bars: 50 µm. **(A-D)** xy images and orthogonal section of 2D monolayers. **(E-H)** Maximal intensity z-projection images of A549 spheroids **(I-L)**, xy images and orthogonal section of 3D spheroids. **(M-P)** Zoomed-in xy images of the top left ¼ area of the spheroids shown in (I-L) with white arrows indicating NPs. Additional images are provided in Figure S2 (for 2D models) and Figure S3 (for 3D models). **(Q)** Normalized mean PL intensity of NPs versus spheroid depth. Curves for individual spheroids are shown in Figure S4.

To investigate the effect of NP size on accumulation and penetration in 3D models, A549 spheroids were incubated with the AuNSs for 24 h. Maximum intensity z-projection images of the spheroids after incubation with different-sized AuNSs suggest a linear increase in NP accumulation with size. Only a few AuNS10 were detected within the spheroids (Figures 2E and S3), whereas AuNS30, AuNS50, and AuNS65 showed progressively higher levels of accumulation (Figure 2F-H and S3). However, in z-projection images, where xy planes are combined across the 3D structure, NPs in the inner region cannot be differentiated from those on the surface. This distinction is instead clear in single xy planes (mid-spheroid) and orthogonal xz and yz, which show a consistent trend: almost no NPs for AuNS10 (Figure 2I), minimal for AuNS30 (Figure 2J), a moderate increase for AuNS50 (Figure 2K), and a significantly higher accumulation for AuNS65 (Figure 2L). Overall, both 2D and 3D data indicate that AuNS10 has the lowest internalization, while AuNS65 shows the highest one among the four sizes tested.

Magnified xy images reveal that NP size also impacts spheroid penetration (Figures 2M-P and S3). Despite their very low number, AuNS10 were evenly distributed between the peripherical layers and the spheroid center (Figure 2M, white arrows) and AuNS30 were mainly located in the first two peripherical cell layers, with some penetrating slightly deeper (Figure 2N, white arrow). In contrast, AuNS50 and AuNS65 showed limited penetration, being mostly confined to the outer layers (Figure 2O-P).

To confirm these observations, the photoluminescence (PL) intensity of AuNSs across spheroid depth was analyzed using custom software. Briefly, each spheroid was divided into 5 μm thick rings and the mean PL per ring was calculated (Figure S13). Since the amount of accumulated particles varied, the PL curves for each spheroid were normalized to highlight differences in penetration depth (Figure 2Q). For all four NPs the PL curves decayed with penetration depth. AuNS10 showed the deepest penetration and an irregular decay, with PL intensity linearly dropping up to ∼30 μm from the surface and slightly increasing at 45 µm, 65 µm, and 85 µm. These variations and high standard deviation (SD) of PL values resulted from the low NP accumulation but also pointed to the presence of NPs at different depths, as shown in the fluorescence images (Figure 2M). AuNS30, AuNS50, and AuNS65 penetration curves decayed very similarly, with the PL drastically dropping at ∼30-40 µm depth, corresponding to about two cell layers. The AuNS30 curve had a slight increase at ∼70 µm, indicating the presence of some NPs in deeper layers, as shown in Figure 2N. In summary, larger NPs accumulated more, while smaller NPs penetrated deeper into the spheroids.

### Influence of the surface charge

To investigate the influence of surface charge on NP cellular uptake, A549 cell monolayers were incubated with three differently charged AuNRs for 3 h. Confocal fluorescence images revealed charge-dependent differences in cellular uptake: AuNR(-) had the highest internalization, followed by AuNRØ, while AuNR(+) showed minimal uptake (Figures 3A-C and S5). The internalization of AuNRØ varied among cells, with some showing NP aggregates.

**Figure 3.**
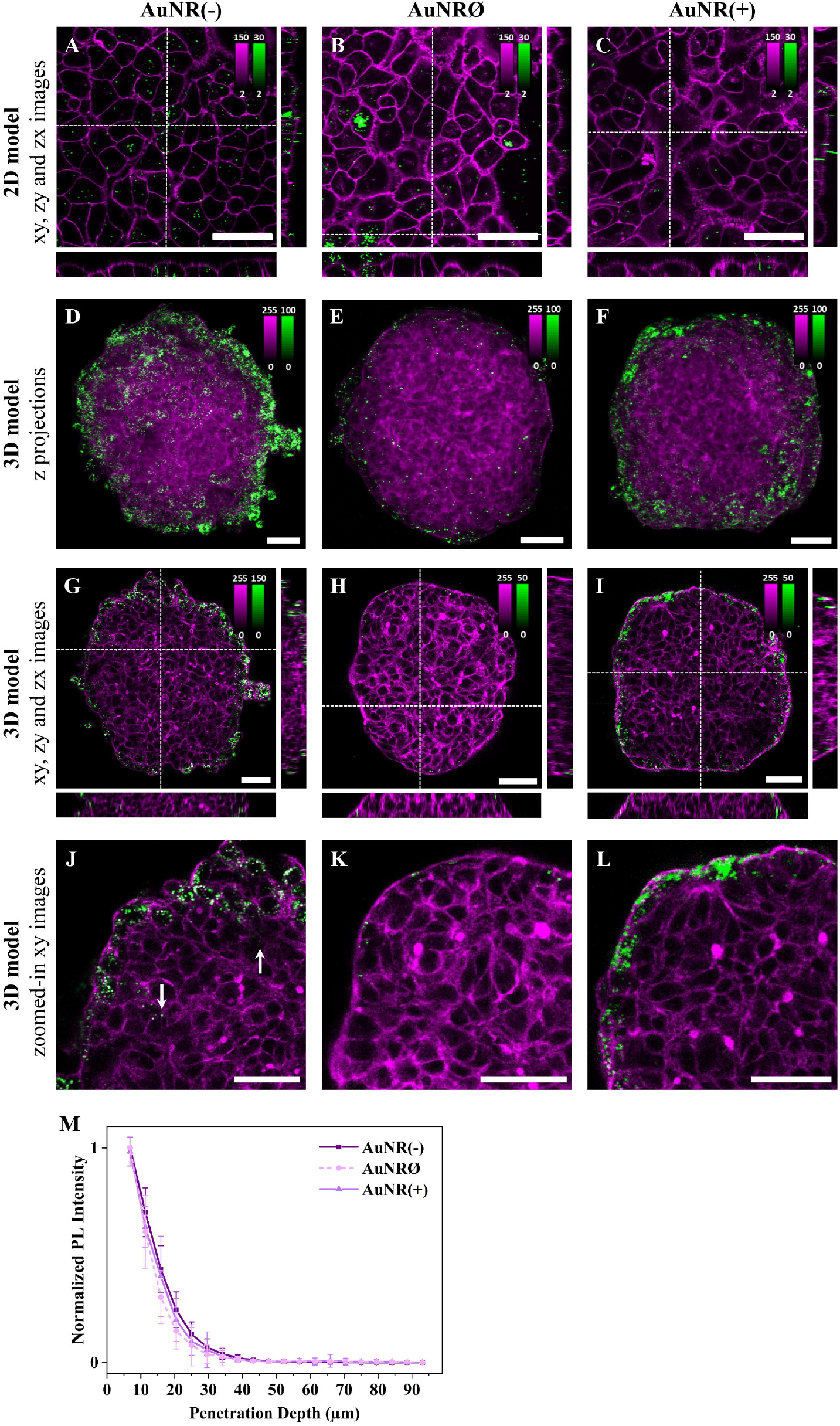
The effect of NP charge on cellular uptake and spheroid penetration. Representative confocal fluorescence images of AuNR(-), AuNRØ, and AuNR(+) in **(A-C)** 2D monolayers and **(D- L)** 3D spheroids of A549 cells. Cells were incubated with ∼4×10^11^ NPs/mL for 3 h or ∼4×10^11^ NPs/mL for 24 h, for 2D or 3D models, respectively. NP PL is shown in green while cellular staining appears in magenta (CellMask^TM^ Green plasma membrane staining for 2D models and Phalloidin CruzFluor^TM^ 488 for cytoskeleton staining in 3D models). Scale bars: 50 µm. **(A-C)** xy images and orthogonal section of 2D monolayers. **(D-F)** Maximal intensity z-projection images of A549 spheroids **(G-I)** xy images and orthogonal section of 3D spheroids. The intensity range is adjusted for each image to highlight NP penetration. Images with the same intensity range are shown in Figure S7. **(J-L)** Zoomed-in xy images of the top left ¼ area of the spheroids reported in (G-I). Additional images are provided in Figure S5 (for 2D models) and Figure S6 (for 3D models). **(M)** Normalized mean PL intensity of NPs versus spheroid depth. Curves for individual spheroids are shown in Figure S7.

To assess the impact of NP surface charge on accumulation and penetration in 3D models, A549 spheroids were incubated with AuNR(-), AuNRØ, or AuNR(+) for 24 h. Maximum intensity z-projections of spheroids, incubated with the differently charged NPs, showed the highest accumulation for AuNR(-), followed by AuNR(+), where a substantial amount of NPs was still detected, and AuNRØ, with a very low number of NPs (Figures 3D-F and S6). Overall, AuNR(-) had the highest internalization in both 2D and 3D models, while for AuNRØ, which had a high uptake in 2D, very few particles were detected in 3D spheroids. The amount of internalized NPs observed in the xy images confirmed this accumulation trend (Figure 3G-I and S6).

Regarding penetration, magnified xy images show that most NPs, regardless of the charge, are confined to the first one or two outer cell layers of the spheroid (Figure 3J-L and S6). Despite a few AuNR(-) particles being found deeper inside the spheroid (Figure 3J, white arrows), the penetration curves revealed no significant differences between the different NPs (Figure 3M), as these inner NPs accounted for a very small fraction of the total particles detected. In summary, negatively charged NPs showed the highest accumulation and marginally deeper penetration in 3D cell models.

### Influence of NP shape

To explore the influence of the shape on NP cellular uptake, rod-shaped and spherical NPs were compared. Negatively charged NPs were selected to eliminate surface charge effects and enhance cellular uptake, as well as spheroid accumulation (Figure 1I). A549 cell monolayers were incubated with AuNS10, AuNS50, or AuNR(-). AuNS10 and AuNS50 were selected as spherical NPs, as their diameters matched the two dimensions of AuNR(-).

Figures 4A-C show representative fluorescence images for the three conditions. In 2D monolayers, internalization of rod-shaped NPs was lower compared to spherical NPs, with a more significant difference observed with AuNS50 (Figure 4A-C and S9). The larger spherical NPs, indeed, showed the highest uptake, as expected from the size comparison experiments (Figure 2). Of note, differences in internalization are evident between Figures 2-3 and Figure 4 due to the different concentrations used. For this set of experiments, the [AuNR(-)] was reduced, whereas the [AuNSs] was increased to match concentrations across the three samples.

**Figure 4.**
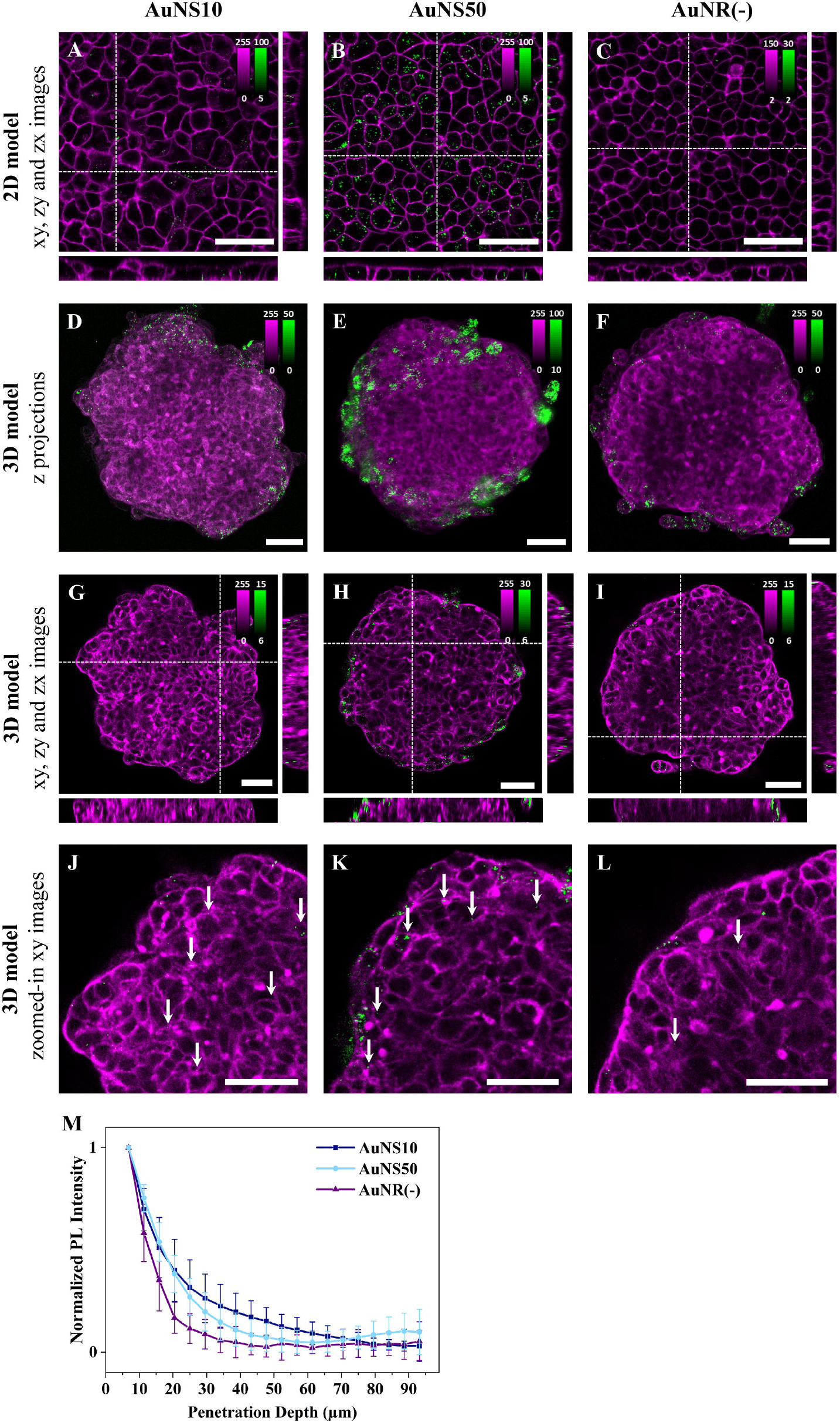
The effect of NP shape on cellular uptake and spheroid penetration. Representative confocal fluorescence images of AuNS10, AuNS50, and AuNR(-) in **(A-C)** 2D monolayers and **(D- L)** 3D spheroids of A549. Cells were incubated with ∼8×10^1^□ NPs/mL for 3 h or ∼5×10^1^□ NPs/mL for 24 h, for 2D or 3D models, respectively. NP PL is shown in green while cellular staining appears in magenta (CellMask^TM^ Deep Red and Green plasma membrane staining for 2D models incubated with AuNSs and AuNRs, respectively, and Phalloidin CruzFluor^TM^ 647 and 488 for cytoskeleton staining in 3D models incubated with AuNSs and AuNRs, respectively). Scale bars: 50 µm. **(A-C)** xy images and orthogonal section of 2D monolayers. **(D-F)** Maximal intensity z-projection images of A549 spheroids **(G-I)** xy images and orthogonal section of 3D spheroids. The intensity range is adjusted for each image to highlight NP penetration. Images with the same intensity range are shown in Figure S11. **(J-L)** Zoomed-in xy images of the top left ¼ area of the spheroids reported in (G-I). Additional images are provided in Figure S9 (for 2D models) and Figure S10 (for 3D models). **(M)** Normalized mean PL intensity of NPs versus spheroid depth. Curves for individual spheroids are shown in Figure S12.

For the experiments in 3D models, A549 spheroids were incubated with AuNS10, AuNS50, and AuNR(-) for 24 h. Based on maximum intensity z-projections and single xy images, NP accumulation follows a trend consistent with the results from the 2D models: AuNR(-) had the lowest accumulation, followed by AuNS10, while AuNS50 exhibited a significantly higher accumulation (Figures 4D-I and S10). Regarding NP penetration, magnified xy images highlight that AuNS10 penetrated deeper into the spheroid (Figure 4J, white arrows). By contrast, AuNS50 primarily accumulated in the first two to three peripherical cell layers (Figure 4K, white arrows), with a small fraction occasionally reaching deeper, close to the spheroid core (Figure S10E, white arrow). AuNR(- ) remained mostly in the first two cell layers (Figure 4L, white arrows). As depicted in Figure 4M, calculated penetration curves confirm these findings. AuNS10 penetration gradually decreased with spheroid depth reaching negligible values around 70 µm. The AuNS50 curve presented a sharp drop at ∼30-40 µm with a rise near 75 µm, indicating the presence of some NPs at the spheroid core. The AuNR(-) curve decreased rapidly at ∼35 µm (about two cell layers), and flattened beyond this depth. Overall, spherical NPs achieved the highest internalization and deeper penetration.

## Discussion

In this systematic study, we investigated the influence of NP size, shape, and surface charge on cellular uptake in 2D monolayers, and accumulation/penetration in 3D spheroids. By isolating each NP property, we provide a comprehensive perspective on NP-cell interactions. A major strength of this work is the direct comparison of the effect of nanoparticle size, surface charge and shape across both 2D and 3D models, using the same cell line and NPs made of the same material for consistency.

### Effect of nanoparticle size

Our results reveal that larger NPs are generally internalized more efficiently than smaller ones. The size-dependent variation in cellular uptake can be attributed to several factors. Smaller NPs, like AuNS10, are more affected by Brownian motion, leading to increased random movement and fewer interactions with the cell membrane, thereby limiting internalization.^7^ In contrast, larger NPs, such as AuNS65, tend to sediment onto the cell membrane, promoting cellular uptake. Membrane wrapping during endocytosis, which is more efficient for larger NPs,^26,27^ may also contribute to the size-dependent NP internalization. NPs between 30-60 nm are more effective at promoting receptor binding, which enhances membrane wrapping and endocytosis. Smaller NPs, like AuNS10, are less effective at initiating this process due to reduced receptor interactions. Despite the trend observed, AuNS50 had a lower internalization than AuNS30. This suggests possible differences in endocytic or exocytosis pathways, though the exact reason remains unclear.

NP behavior in 3D models is shaped by factors beyond uptake mechanisms, including ECM density and diffusion barriers. Smaller NPs like AuNS10 penetrate deeper into spheroids, likely due to increased Brownian motion, which reduces NP internalization but improves diffusion through the ECM and interstitial spaces.^28^ Larger NPs, however, experience stronger NP-cell interactions and increased cellular uptake, but limited diffusion through the ECM, which confines them to the peripheral cell layers. These results show that while larger NPs accumulate efficiently in solid tumors, they lack the penetration depth achieved by smaller NPs. These findings highlight the need for designing the next-generation NPs with smaller sizes and bioactive surfaces to enhance internalization through targeted interactions.^29^

### Effect of surface charge

Our study demonstrates that negatively charged NPs exhibit the highest internalization, followed by neutral and positive NPs. Although positive NPs are generally expected to interact with the negatively charged plasma membrane,^30^ enhancing NP internalization,^31^ AuNR(+) showed the lowest uptake in 2D monolayers. This unexpected contradiction can be explained by the change of the surface charge of AuNR(+) in cell culture medium, as indicated by a drop in the Zeta potential (from 57 to -13 mV, Figure 1I). In the medium, a protein corona, mainly composed of negatively charged proteins like albumin,^32^ can form around the positively charged NPs. This protein corona acts as a steric barrier, reducing close contact with the plasma membrane and, in turn, the likelihood of endocytosis.^33–35^ This result showcases the importance of evaluating surface charge in culture medium. The increased internalization of AuNRØ in 2D monolayers can be attributed to NP aggregation, which occurs more easily due to their lower colloidal stability.^36,37^ NP aggregates are more readily taken up by the cells, as their larger size promotes sedimentation and increased interactions with the cell membrane.^7^ In spheroids, however, no large aggregates were detected and AuNRØ showed the lowest accumulation.

Analysis of spheroid penetration revealed similar penetration curves for AuNR(-), AuNRØ, and AuNR(+). However, confocal microscopy images consistently showed that a small fraction of AuNR(-) reached deeper layers compared to the other NPs. This enhanced penetration of AuNR(-) is likely due to electrostatic repulsion with the negatively charged ECM, which facilitates diffusion.^38^ Although AuNR(+) acquired a negative charge in the medium, its protein corona likely restricted its movement, keeping it concentrated at the spheroid’s periphery.^38^ AuNRØ, while preventing strong interactions, are hindered by their tendency to form bigger aggregates causing steric hindrance within the tumor structure. These findings underscore the efficient internalization and superior spheroid penetration of negatively charged NPs and spotlight the importance of surface charge in the design of NPs for cancer therapy.

### Effect of nanoparticle shape

The shape of NPs affects their internalization in both 2D and 3D models. In our study, spherical AuNS50 and AuNS10 exhibit higher uptake compared to rod-shaped AuNR(-) in both 2D and 3D models. This difference is likely due to the higher energy barrier faced by rod-shaped NPs for cellular internalization, as membrane wrapping is less efficient.^39^ Additionally, rod-shaped NPs can enter cells at various angles, which also impacts their uptake rate.^39,40^

In the spheroid model, spherical NPs showed deeper penetration than rod-shaped NPs, probably due to differences in interactions of each shape with the dense ECM network.^15^ Rod-shaped NPs encounter greater steric hindrance due to their elongated form, making it harder to navigate through tight spaces and limiting their diffusion in spheroids. Spherical NPs, on the other hand, reach deeper layers, with the occasional detection of AuNS50 in the spheroid core. Of note, when lower concentration was used, this core fraction was not detected due to reduced NP accumulation (Figure 2).

In conclusion, this research highlights the value of 3D spheroid models in better understanding NP interactions, offering insights not visible in 2D monolayers. Our results revealed key differences between the two models: while 2D models provide a preliminary understanding of NP internalization, they often do not correlate with *in vivo* behaviour. Spheroids, on the other hand, enabled us to observe the interplay between NP and cells within the tumor, which drives internalization, and NP-ECM interactions, which govern deeper penetration. By leveraging advanced 3D *in vitro* models, researchers can achieve a more comprehensive characterization of NP behaviour, providing more reliable data that can bridge the gap between preclinical findings and clinical outcomes. This approach holds promise for developing optimized nanomaterials with improved performance and a higher likelihood of clinical success in cancer therapy.

## Acknowledgments

This project has received funding from the European Union’s Horizon 2020 research and innovation program under the Marie Skłodowska-Curie grant agreement no. 860914. We acknowledge additional financial support from Research Foundation of Flanders (FWO) research grants (G0D4519N, G081916N, VS08523N, G0C1821N, G022724N), postdoctoral fellowships (for BF: 12X1419N and 12X1423N, for IVZ: 12A6N25N), and from the KU Leuven (IDN/20/021, C14/15/053, C14/19/079, C14/22/085). MB acknowledges the Global PhD partnership program between KU Leuven and Melbourne university (GPUM/21/025).

