## Supplementary material for "Nanoparticle Accumulation and Penetration in 3D Tumor Models: the Effect of Size, Shape, and Surface Charge": Figure S

### 1 Supplementary Figures

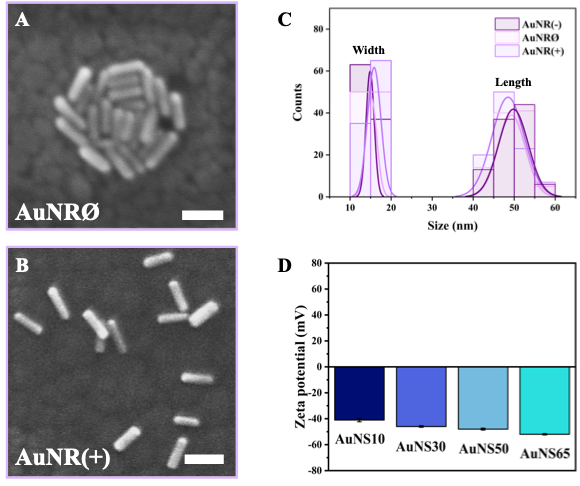

**Figure S1.** Nanoparticle characterization. TEM images of AuNRØ (A) and AuNR(+) (B) (scale bars: 50 nm). Size distribution of AuNRs (C). Zeta potential of AuNSs (D).

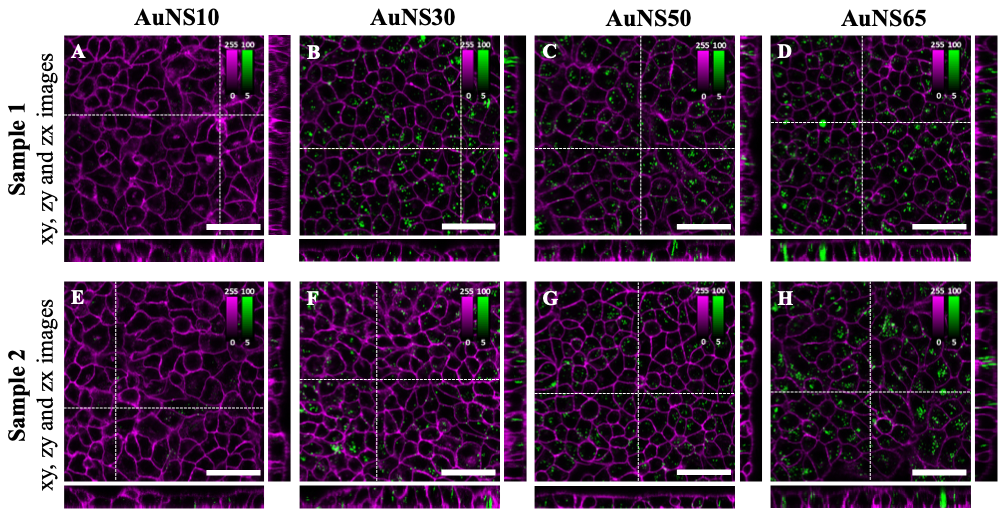

**Figure S2.** The effect of NP size on cellular uptake. Confocal fluorescence imaging of AuNS10 (A,E), AuNS30 (B,F), AuNS50 (C,G), and AuNS65 (D,H) in 2D monolayers of A549 cells after 3 h of incubation. NP PL is shown in green while cells appear in magenta (CellMask^TM^ Deep Red plasma membrane staining). The central panel displays an xy plane within the cells, while the right and bottom panels show the yz and xz orthogonal views. Two different samples are represented for each condition (A-D and E-H, respectively).

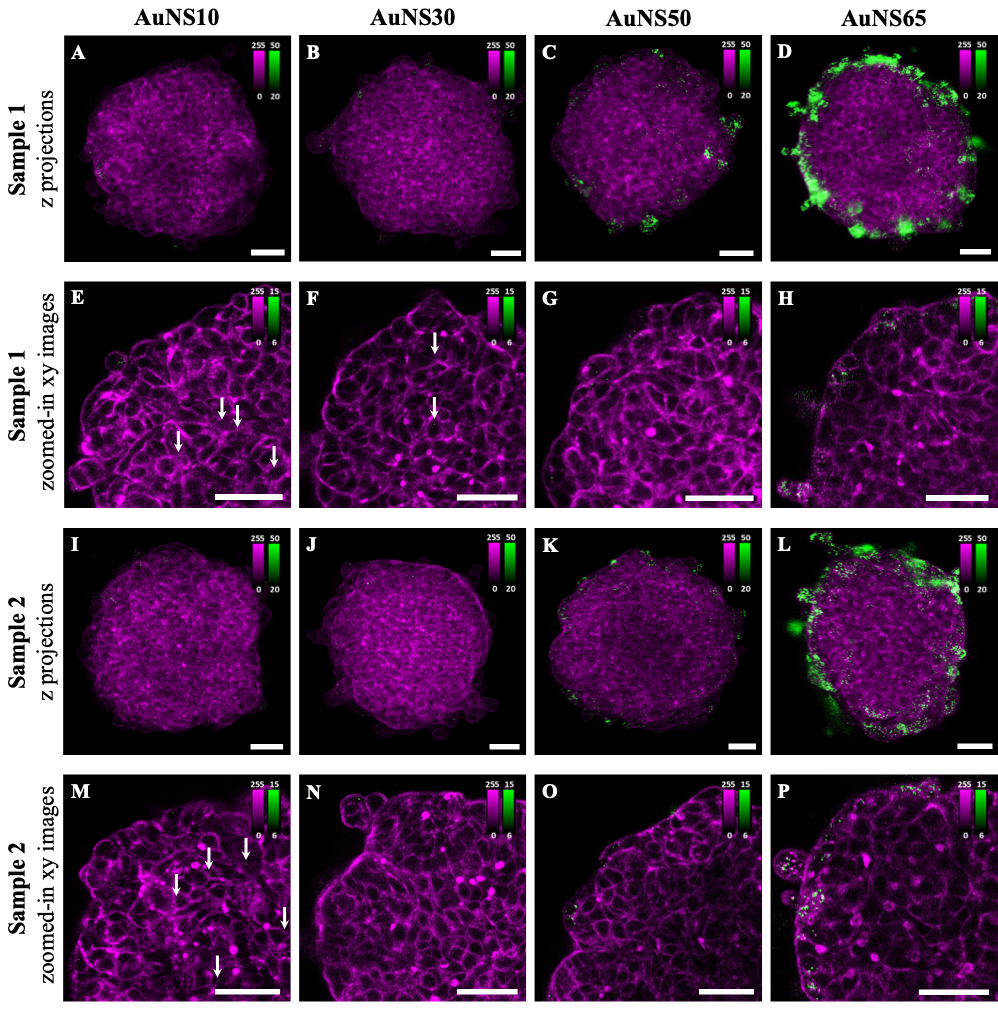

**Figure S3.** The effect of NP size on spheroid accumulation and penetration. Confocal fluorescence imaging of AuNS10, AuNS30, AuNS50, and AuNS65 in 3D spheroids of A549 after 24 h of incubation. NP PL is shown in green while cells appear in magenta (Phalloidin CruzFluor^TM^ 647 for cytoskeleton staining). (A-D,I-L) Z projections of AuNS10, AuNS30, AuNS50, and AuNS65 in A549 3D spheroids. (E-H,M-P) Zoomed-in xy images of the top left ¼ area of the spheroids reported in (A-D,I-L) with white arrows indicating NPs. Two different samples are represented for each condition (A-H and I-P, respectively).

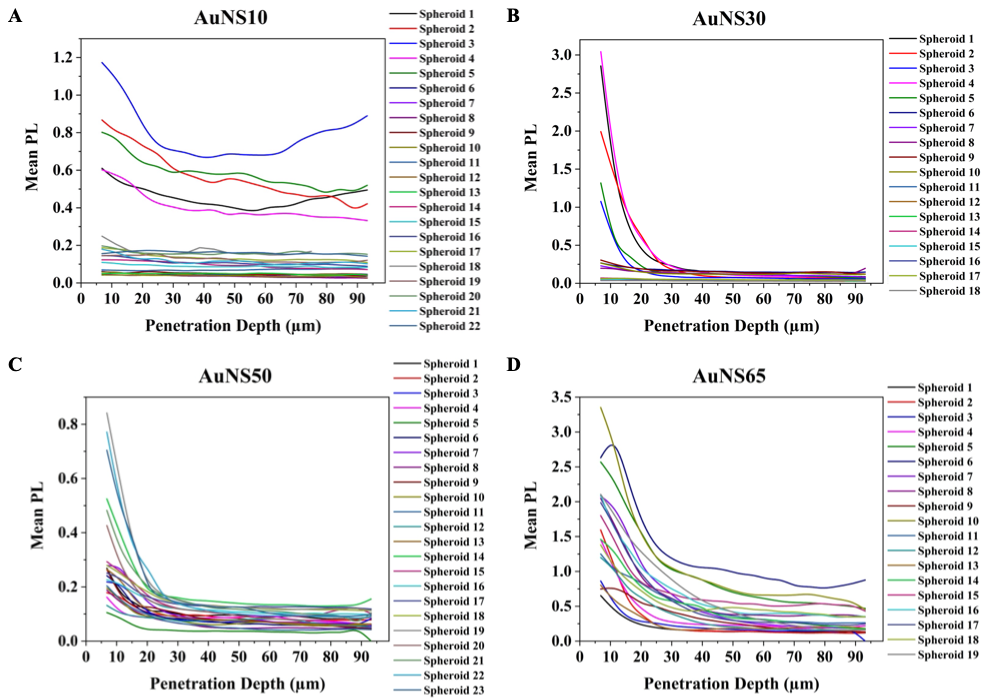
**Figure S4.** The effect of NP size on spheroid penetration. Raw PL intensity of NPs versus spheroid depth represented for AuNS10 (A), AuNS30 (B), AuNS50 (C), AuNS65 (D). Each curve represents the penetration of NPs in a different spheroid.

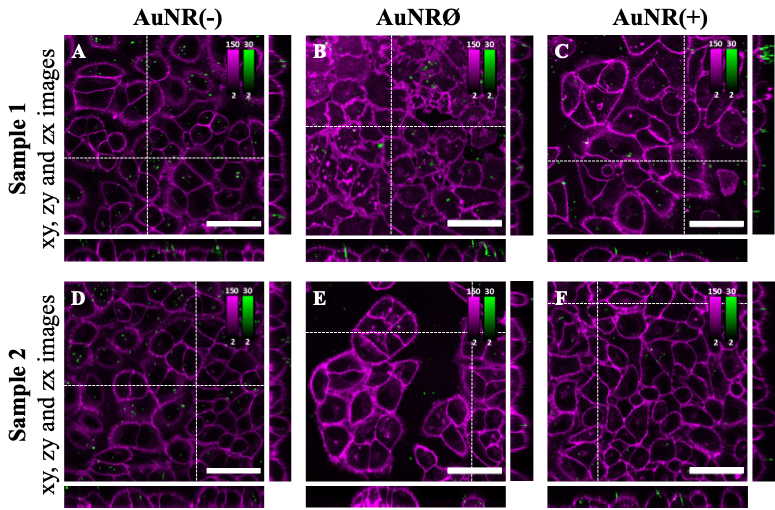

**Figure S5.** The effect of NP charge on cellular uptake. Confocal fluorescence imaging of AuNR(-) (A,D), AuNRØ (B,E), AuNR(+) (C,F) in 2D monolayers of A549 cells after 3 h of incubation. NP PL is shown in green while cells appear in magenta (CellMask^TM^ Green plasma membrane staining). The central panel displays an xy plane within the cells, while the right and bottom panels show the yz and xz orthogonal views. Two different samples are represented for each condition (A-C and D-F, respectively).

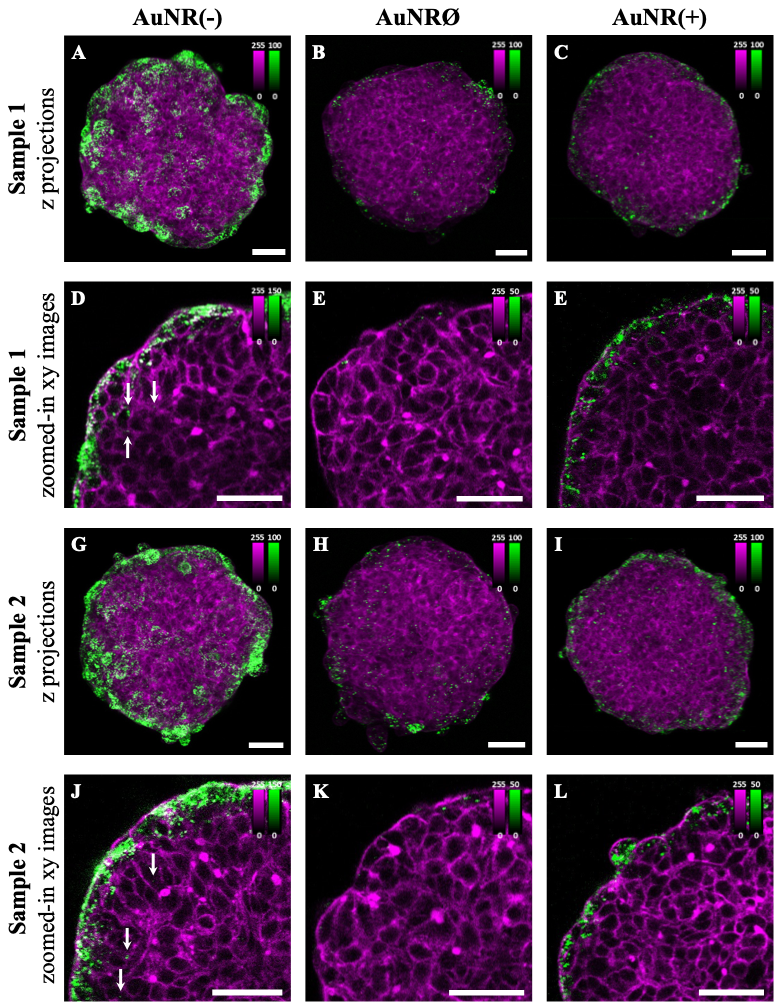

**Figure S6.** The effect of NP charge on spheroid accumulation and penetration. Confocal fluorescence imaging of AuNR(-), AuNRØ, and AuNR(+) in 3D spheroids of A549 after 24 h of incubation. NP PL is shown in green while cells appear in magenta (Phalloidin CruzFluor^TM^ 488 for cytoskeleton staining). (A-C,G-I) Z projections of AuNR(-), AuNRØ, and AuNR(+) in A549 3D spheroids. (D-E,J-L) Zoomed-in xy images of the top left ¼ area of the spheroids reported in (A-C,G-I) with white arrows indicating NPs. Two different samples are represented for each condition (A-E and G-L, respectively).

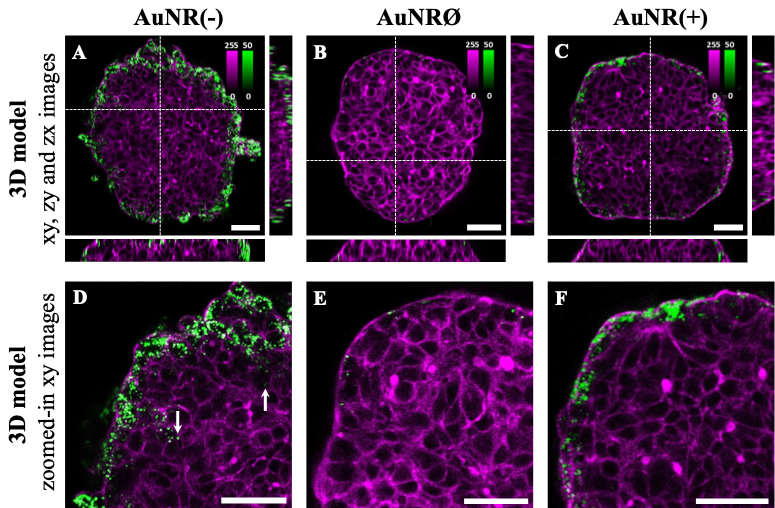

**Figure S7.** The effect of NP charge on spheroid accumulation and penetration. Images from Figure 3 with identical color bars across all conditions.

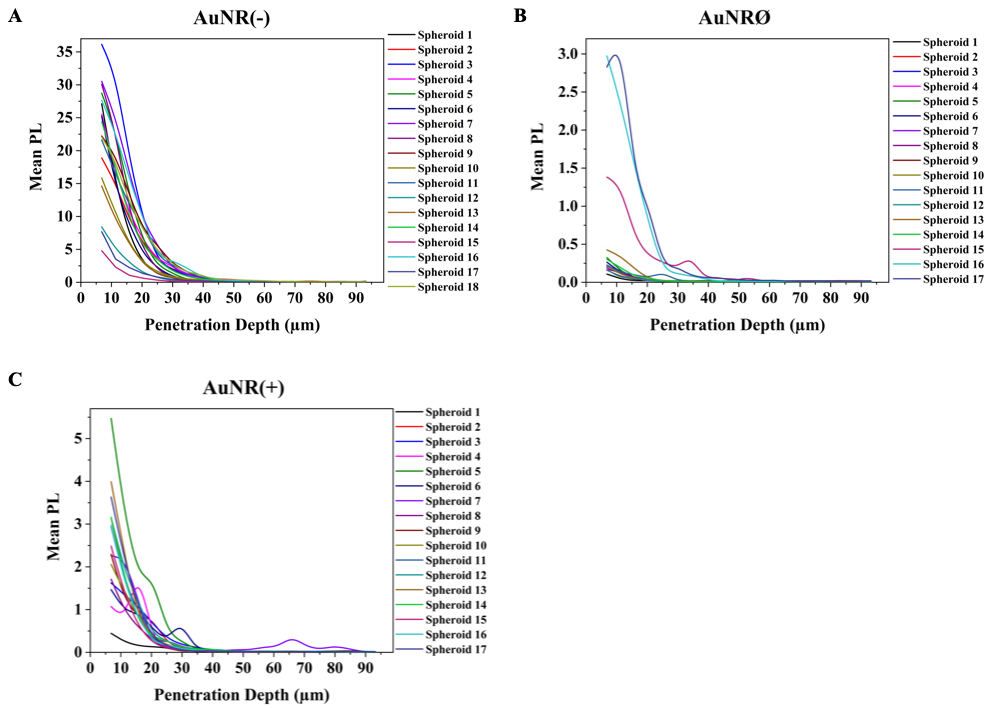

**Figure S8.** The effect of NP charge on spheroid penetration. Raw PL intensity of NPs versus spheroid depth represented for AuNR(-) (A), AuNRØ (B), and AuNR(+) (C). Each curve represents the penetration of NPs in a different spheroid.

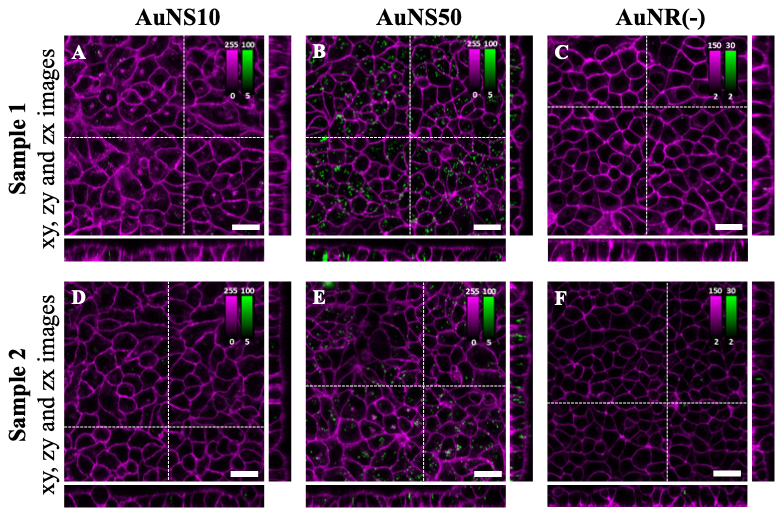

**Figure S9.** The effect of NP shape on cellular uptake. Confocal fluorescence imaging of AuNS10 (A,D), AuNS50 (B,E), AuNR(-) (C,F) in 2D monolayers of A549 cells after 3 h of incubation. NP PL is shown in green while cells appear in magenta (CellMask^TM^ Deep Red plasma membrane staining for AuNSs and CellMask^TM^ Green for AuNR(-)). The central panel displays an xy plane within the cells, while the right and bottom panels show the yz and xz orthogonal views. Two different samples are represented for each condition (A-C and D-F, respectively).

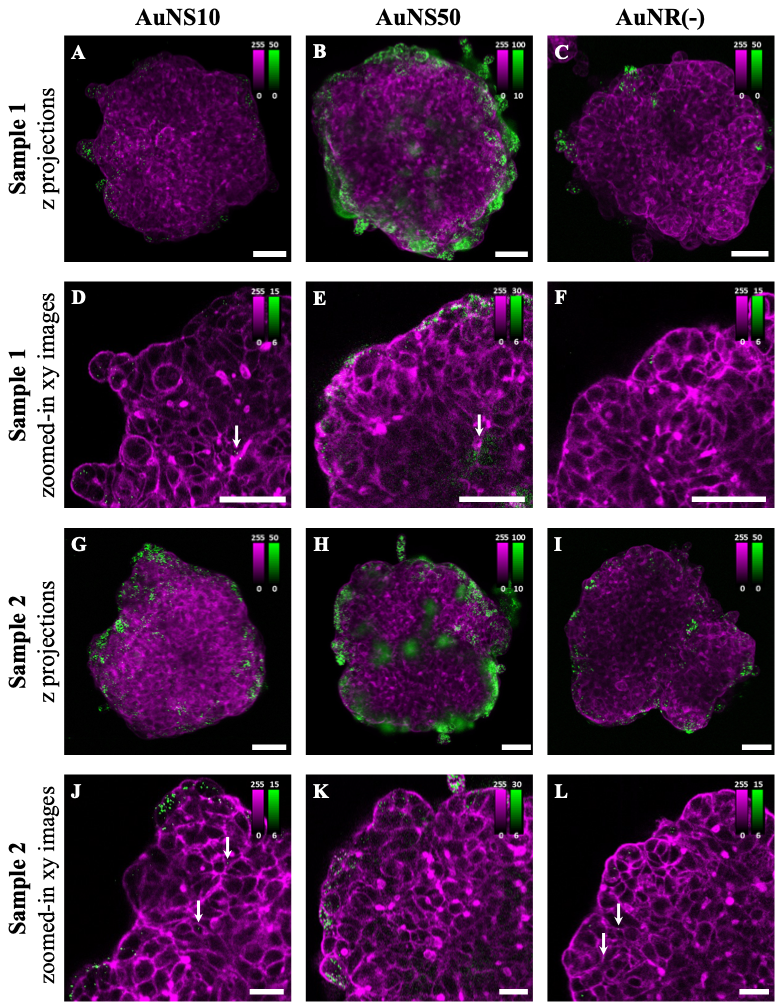

**Figure S10.** The effect of NP shape on spheroid accumulation and penetration. Confocal fluorescence imaging of AuNR(-), AuNRØ, and AuNR(+) in 3D spheroids of A549 after 24 h of incubation. NP PL is shown in green while cells appear in magenta (Phalloidin CruzFluor^TM^ 647 for cytoskeleton staining for AuNSs and Phalloidin CruzFluor^TM^ 488 for AuNR(-)). (A-C,G-I) Z projections of AuNS10, AuNS50, and AuNR(-) in A549 3D spheroids. (D-F,J-L) Zoomed-in xy images of the top left ¼ area of the spheroids reported in (A-C,G-I) with white arrows indicating NPs. Two different samples are represented for each condition (A-F and G-L, respectively).

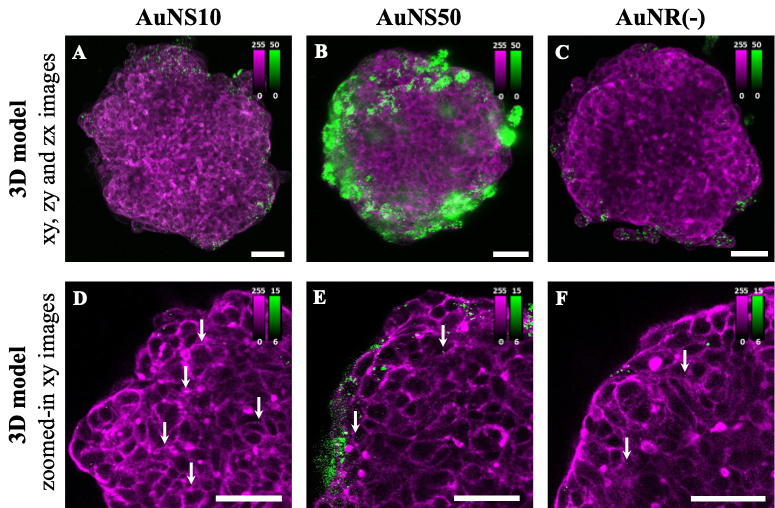

**Figure S11.** The effect of NP shape on spheroid accumulation and penetration. Images from Figure 4 with identical color bars across all conditions.

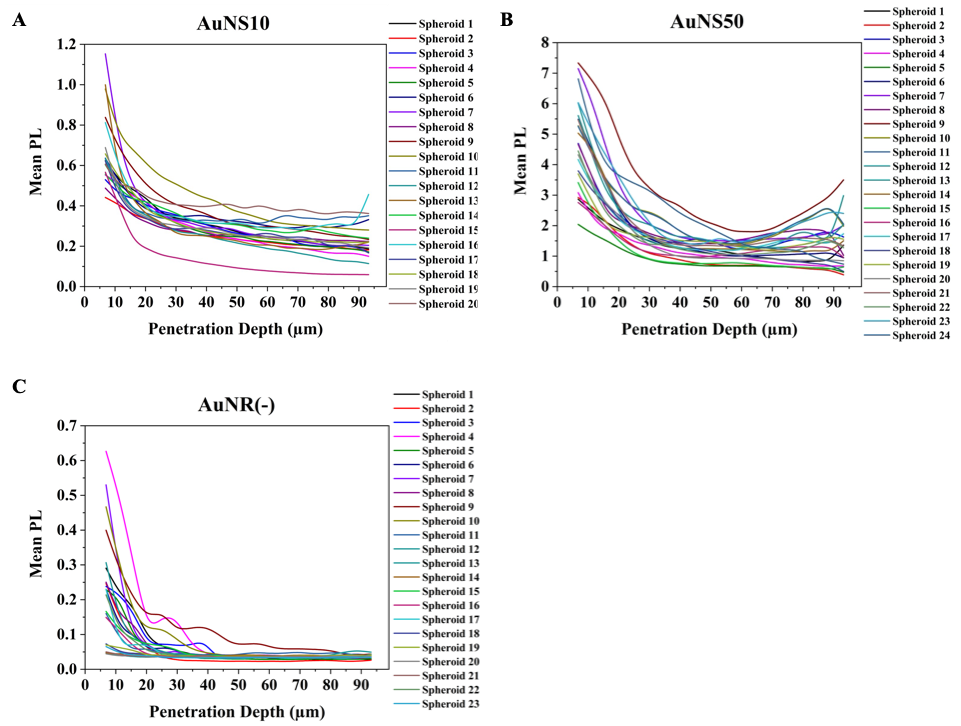

**Figure S12.** The effect of NP charge on spheroid penetration. Raw PL intensity of NPs versus spheroid depth represented for AuNS10 (A), AuNS50 (B), and AuNR(-) (C). Each curve represents the penetration of NPs in a different spheroid.

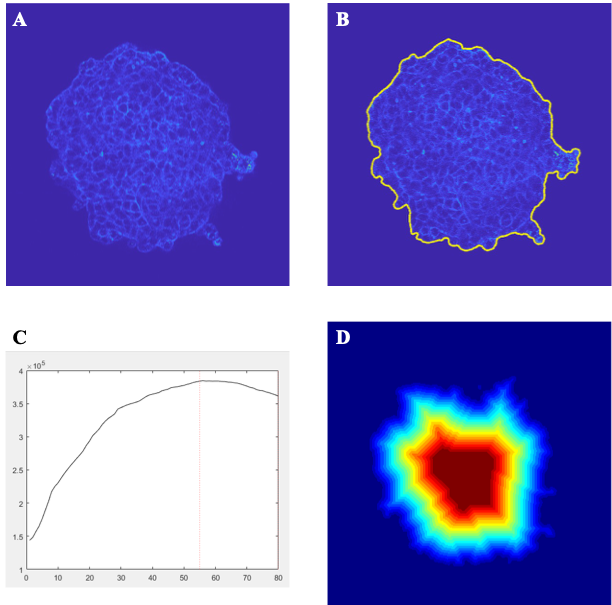

**Figure S13.** Overview of functioning software calculating the mean NP PL intensity over the spheroid depth. (A) xy plane of the spheroid where the user is asked to select an area corresponding to the background and an area corresponding to the fluorescence signal. (B) Detection of the spheroid by the software for each xy plane of the z stack. (C) Evolution of the spheroid diameter over the spheroid depth used to select the z stacks to analyze. (D) Segmentation of each xy plane into rings of the identical inputted width followed by the calculation of the mean NP PL in each ring.
